# Model of a bilateral Brown-type central pattern generator for symmetric and asymmetric locomotion

**DOI:** 10.1101/146993

**Authors:** Anton Sobinov, Sergiy Yakovenko

## Abstract

The coordinated activity of muscles is produced in part by spinal rhythmogenic neural circuits, termed central pattern generators (CPGs). A classical CPG model is a system of coupled oscillators that transform locomotor drive into coordinated and gait-specific patterns of muscle recruitment. The network properties of this conceptual model can be simulated by a system of ordinary differential equations with a physiologically-inspired coupling locus of interactions capturing the timing relationship for bilateral coordination of limbs in locomotion. While most similar models are solved numerically, it is intriguing to have a full analytical description of this plausible CPG architecture to illuminate the functionality within this structure and to expand it to include steering control. Here, we provided a closed-form analytical solution contrasted against the previous numerical method. The computational load of the analytical solution was decreased by order of magnitude when compared to the numerical approach (relative errors, <0.01%). The analytical solution tested and supported the previous finding that the input to the model can be expressed in units of the desired limb locomotor speed. Furthermore, we performed parametric sensitivity analysis in the context of controlling steering and documented two possible mechanisms associated with either an external drive or intrinsic CPG parameters. The results identify specific propriospinal pathways that may be associated with adaptations within the CPG structure. The model offered several network configurations that may generate the same behavioral outcomes.

**New & Noteworthy:** Using a simple process of leaky integration, we developed an analytical solution to a robust model of spinal pattern generation. We analyzed the ability of this neural element to exert locomotor control of the signal associated with limb speeds and tested the ability of this simple structure to embed steering control using the velocity signal in the model’s inputs or within the internal connectivity of its elements.

## Introduction

Specialized neural elements in the spinal cord, known as the central pattern generators (CPGs), contribute to the generation of periodic coordinated patterns of locomotor activity (Grillner and Zangger, 1975). Discovered in deafferented preparations, CPGs do not require sensory signals to produce locomotor behavior; however, their pattern is greatly influenced by sensory and descending inputs (Yakovenko, 2011; Prochazka and Ellaway, 2012). Specifically, the direct electrical stimulation of a brainstem structure called the mesencephalic locomotor region (MLR), even in decerebrated animals, produces oscillations in the CPGs and subsequent locomotor behavior (Grillner and Wallén, 1985). This locomotor behavior is characterized by the complex coordinated actions of multiple muscle groups. It is remarkable that a change in either the magnitude or frequency of MLR stimulation can generate all appropriate modifications of these patterns. This increase in stimulation expresses a full repertoire of gaits with continuous transitions, such as from walking to trotting or galloping in over-the-ground locomotion (Shik et al., 1966), or transitioning from slow walking to swimming in amphibians (Cabelguen et al., 2003), which is faster than walking mode of locomotion. Thus, increasing stimulation input current corresponds to an increase in locomotor velocity.

Many CPG models were developed over the last century (Verzár, 1923; Taga et al., 1991; Bashor, 1998; Yakovenko et al., 2005; Rybak et al., 2006; Markin et al., 2010; Barnett and Cymbalyuk, 2014a). Simulated model structure and its parameters are usually derived from observing the motor output patterns or their changes in response to external inputs or naturally occurring variations. These models give rise to the mechanistic descriptions that capture biological organization and the processes; however, they generally start as phenomenological or statistical representations of observed phase variations or timing in the recorded muscle activity. For example, both the limb-based Brown’s CPG (Brown, 1911) and the joint-based Grillner’s CPG (Grillner, 1981) are similarly founded on the observations of multiple representative electromyographic (EMG) profiles providing insight into the functional organization of this circuitry.

The idea of a CPG as a distributed mechanism that integrates convergent inputs (Grillner and Wallén, 1985) has been supported by both computational and experimental studies. Using calcium imaging, the spatiotemporal activity of rhythmogenic circuitry was found to be functionally distributed with motoneurons in the rostral lumbar and sacral segments of the spinal cord (Bonnot et al., 2002; O’Donovan et al., 2005). The spatiotemporal distribution of neural activity throughout the lumbar enlargement with descending control and sensory inputs intact was visualized by combining the anatomical location of the motoneurons with information about their activity during normal locomotion (Yakovenko et al., 2002). This was also supported by observations of independent and coupled recruitment of flexor and extensor rhythmogenic spinal circuits using selective optogenetic approaches (Hägglund et al., 2013). The rhythmogenesis in only flexors or only extensors observed with optogenetics supports the computational observation of a switch-like transition between flexors and extensors (or more precisely, limb protractors and retractors), which identifies them as distinct network elements (Yakovenko et al., 2002). This bilateral, switch-like activation of the motor pools spanning the full rostocaudal extent of the lumbosacral enlargement is likely associated with distributed rhythm-generating networks responsible for this activity.

The integration of feedforward predictions and sensory feedback about ongoing execution is the optimal solution for generating robust control of complex body morphology (Kuo, 2002). Over the course of evolution, the process of optimization within control pathways has likely been concerned with the optimization of locomotion, as this is a central behavior that is essential for animal survival (Yakovenko, 2011). One engineering solution to the problem of computing predictive commands for complex systems is the use of inverse models (Smith, 1957; Wolpert and Ghahramani, 2000). The complex transformation from muscle activations into movement kinematics could be internalized for inverse solutions that generate appropriate output for the desired kinematic input. It is then not surprising that dedicated rhythmogenic networks for locomotion may be embedding the dynamics of body-ground interactions to solve the problems of intra- and interlimb coordination (Taga et al., 1991; Full and Koditschek, 1999). The accuracy of these embedded neural calculations of musculoskeletal transformation may be fine-tuned by experience (Wolpert et al., 1998; Bhushan and Shadmehr, 1999; Kawato, 1999; Ijspeert et al., 2013). It is important to acknowledge that sensory feedback pathways may also shape the final output of motor pathways and compensate for dynamics during locomotion. In addition, there is considerable evidence that CPGs integrate sensory inputs together with supraspinal commands to generate changes in the timing and magnitude of locomotor activity (Ijspeert, 2008; Yakovenko, 2011).

CPG models offer a unique research opportunity to understand the interplay between these neural directives and biomechanical constraints that govern a complex dynamic task. To this extent, we have previously used inverse solutions of a CPG model to infer the nature of descending inputs (Yakovenko, 2011). The surprising result of these simulations was that the input to the CPG was the velocity of each limb. Described mathematically as a system of differential equations (Matsuoka, 1985; Schöner et al., 1990; Wallén et al., 1992; Cymbalyuk et al., 2002; Rybak et al., 2006; Yakovenko, 2011), CPG models are hard, even impossible, to solve analytically in the form of known functions and variables. Still, analytical expressions have several advantages over numerical models. Unlike numerical solutions that often suffer from the accumulating errors and inversely related computational load, the analytical solutions are precise within assumptions taken during their derivation. Even though they are also evaluated, their formulation is more efficient and faster than the approximate numerical solutions.

In this study, we developed a method to obtain an analytical solution to one of the simplest implementations of a locomotor CPG. We used this analytical expression to further test the ability of this circuitry to embed the regulation of phases appropriate for different speeds and control steering with asymmetric gaits. While the identification of pattern generating elements is a considerable challenge in experimental techniques, the function of distributed elements of a CPG can be probed with computational methods that allow us to monitor and manipulate any part of the circuit. We tested two hypotheses in this study: 1) the exact analytical solution exists for a bilateral CPG model implemented with a leaky integration process; 2) the intrinsic circuit redundancy in a CPG can accommodate the expression of asymmetric gait. The function of embedding the asymmetric representations of gait may be relevant for understanding steering and short- and long-term adaptations within spinal systems.

## Methods

### Model description

While a few CPG models of neural activity consider specific ion dynamics using the Hodgkin-Huxley formulation (Cymbalyuk et al., 2002; Rybak et al., 2006), our model captures gross CPG network dynamics, described by T.G. Brown, in a form of gated leaky integration. We expressed the input-output relationship using coupled leaky integrators formulated as a system of ordinary differential equations (ODEs). The system of ODEs can be expressed in matrix form (Eq. 1), with ipsilateral antagonism expressed as abrupt, non-overlapping state transitions. An event associated with any given state *x_i_* value crossing 1 triggers the resetting of the state to 0 and the start of integration for the ipsilateral antagonist. In Fig. 1, for example, if the left flexor (*x_1_*) reaches 1, it resets to 0 and turns off, while the left extensor (*x_2_*) switches on.

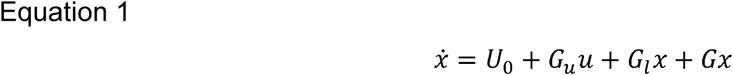

where *x* = *(x_1_, x_2_, x_3_, x_4_)^T^* - state vector, *U_0_* - constant input from intrinsic connections, *G_u_* - extrinsic input gains, *u* - extrinsic inputs, *G_l_* - leak gains, *G* - weights for connections between integrators (*r_ff_, r_fe_, r_ef_, r_ee_* weights in Fig. 1).

**Fig. 1.**
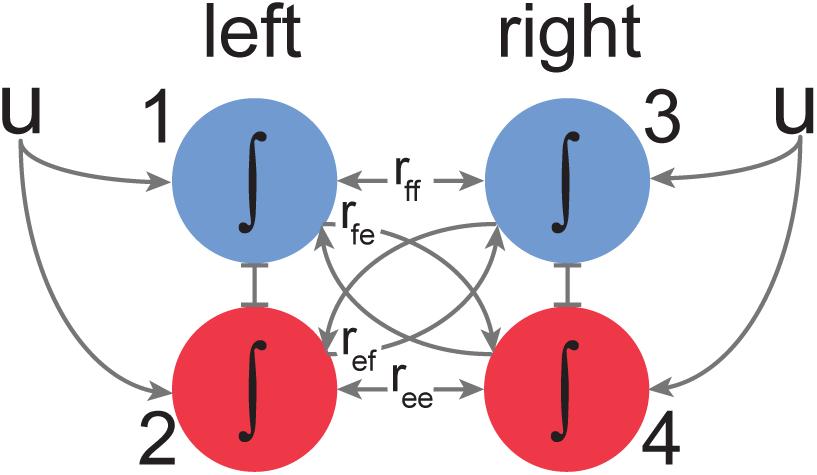
Schematic of the bilateral locomotor CPG model. The oscillatory behavior in each half-center (marked 1-4) was generated through an intrinsic, leaky integrate-to-threshold resetting. This process was also under regulation from intrinsic inputs governed by parameters (*r_ff_, r_fe_, r_ef_, r_ee_*). The flexor half-centers (blue) were reciprocally connected to extensor half-centers (red). See Eq. 1-2 for details.

To simplify model parameter space, the parameters were coupled assuming symmetrical organization across the midline, as seen in Eq. 2. Additionally, the connection between flexors (*r_ff_*) was removed for simulations of walking behavior, where swing phases do not overlap.

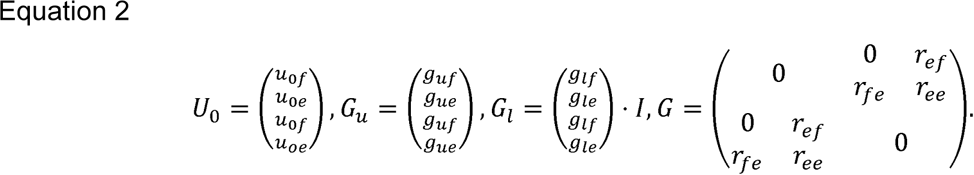

We used the fixed-step 4th order Runge-Kutta method with 10^-3^ s precision for forward numerical integration.

### Analytical solution

The bilateral CPG model produces flexor (swing) and extensor (stance) phases for two limbs in relation to extrinsic input and intrinsic structure. To obtain these phases, Eq. 1 needs to be integrated in time between the state changes. Numerical integration was previously used (Yakovenko, 2011) to generate swing and stance periods. The same transition points can be calculated analytically by transforming Eq. 1 into a matrix Cauchy problem:

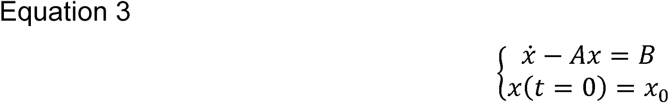

where *A*=*G_l_*+*G* represents the intrinsic structure of the CPG, *B*=*U_0_*+*G_u_u* represents the state-independent inputs, and *x_0_* is the initial condition. In the case of a non-singular matrix *A*, this system has a vector form solution:

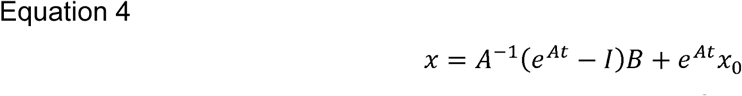

where *I* is the identity matrix. This analytical expression of states *x* (with dimensionality [4×1] for a model of bilateral CPG) describes the progression of all locomotor phases in time between the state changes. The remaining task is then to calculate the transition times and corresponding phase durations for a full step cycle. Eq. 4 was evaluated for all three possible bilateral combinations of concurrent flexor-extensor activity during a full step cycle, namely: i) left flexion and right extension (states *x_1_* and *x_4_*), ii) left extension and right extension (states *x_2_* and *x_4_*), and iii) left extension and right flexion (states *x_2_* and *x_3_*). States may have repeated more than once within the step cycle, when CPG activity was highly asymmetric. The dimensionality of the problem can be reduced from 4 to 2 because only two integrators are active at any given time with the following parameters:

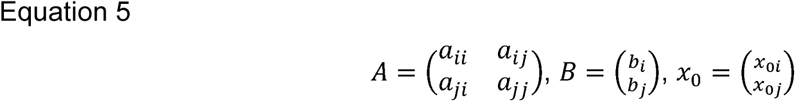

where *i* ⎕ {1,2} and *j* ⎕ {3,4} are the indices of the two active integrators. We can then find the time of phase transitions *τ* for a given integrator *k* by inserting the reduced parameter set (Eq. 5) into Eq. 4 and assuming *x_i_* or *x_j_* is equal to 1. Solving for *τ* yields the following transcendental equation:

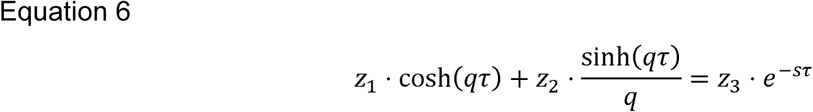

where *z_1_*, *z_2_*, *z_3_*, *s*, *q* are parameters describing the model configuration. τ was then found numerically using Brent’s method and analytically by expanding the hyperbolic functions using a Maclaurin series. We used the NumPy ‘roots’ function (Horn and Johnson, 1999) to solve the polynomials of power over 2. Next, the periods of activity of flexors and extensors during a step cycle were obtained with the following iterative algorithm:

i. Calculate the time *τ_i_* when state *x_i_* reaches 1.
ii. Calculate the time *τ_j_* when state *x_j_* reaches 1.
iii. Calculate the state of all integrators at time point *τ=min(τ_i_, τ_j_)*.
iv. Reset the state from 1 to 0, deactivate it, and activate the reciprocal ipsilateral state. For example, switch from an active left flexor to an active left extensor.
v. If a full step cycle is completed (all 4 states reached value 1 at least once), stop; otherwise, go to step (i).

### Cost function

The CPG model can generate multiple locomotor behaviors as a function of extrinsic inputs and intrinsic interactions (Yakovenko, 2011). Given a desired behavior, e.g. stereotypical symmetrical walking (Halbertsma, 1983), the appropriate CPG parameters were found by optimizing the cost function (Eq. 7) that expressed the goodness of fit between target (experimental) and simulated patterns. In the symmetrical model, we optimized for 6 different speeds, from 0.1 to 1.5 m/s (dashed lines in Fig. 2), that were generated with 6 values of *u* (evenly distributed between 0.1 and 1.5 au). Fig. 2 shows the quality of simulated solutions for symmetrical walking over a full range of walking speeds.

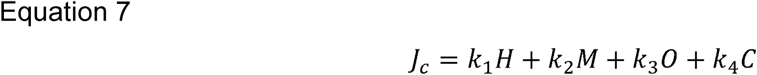

**Fig. 2.**
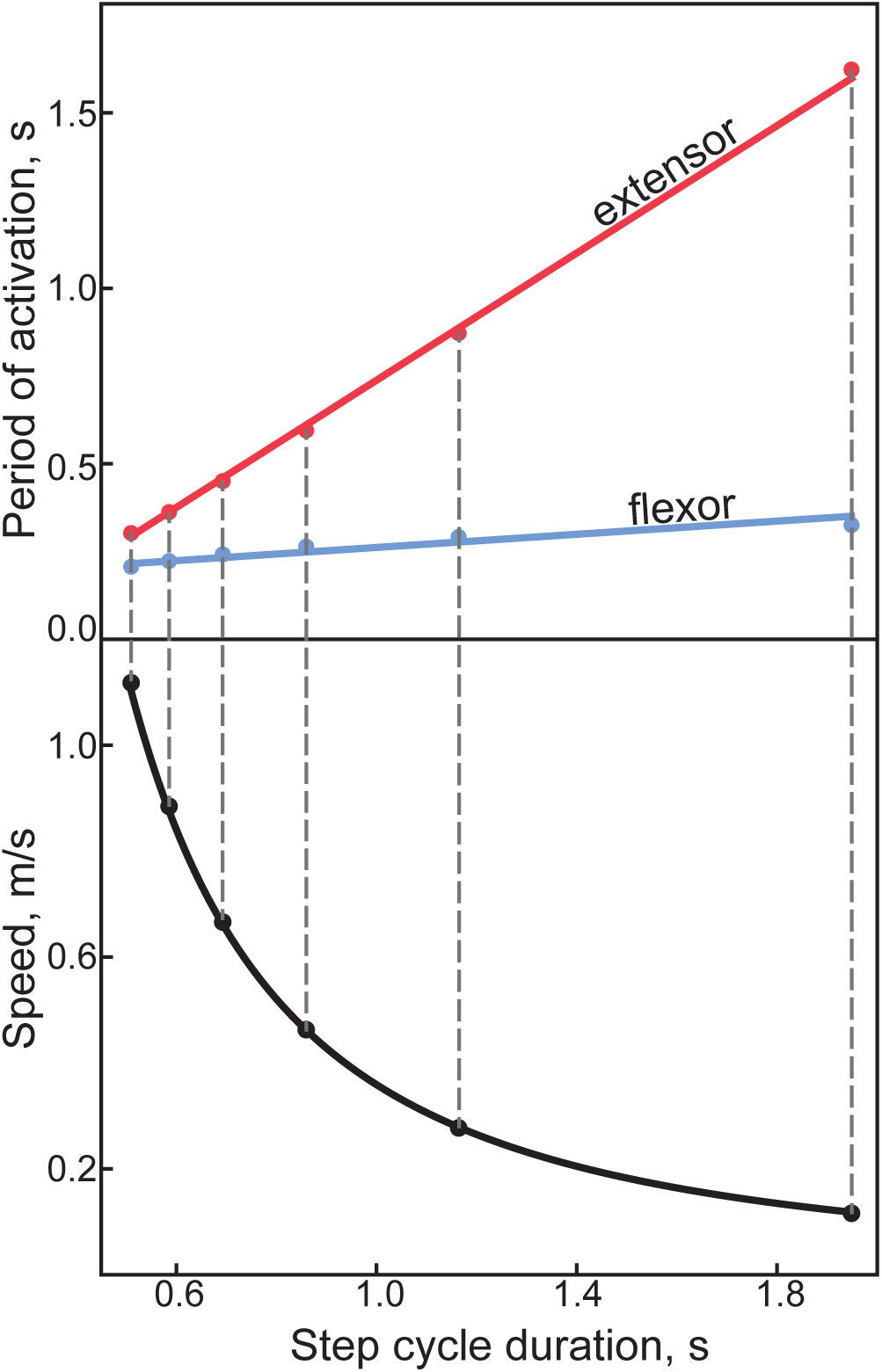
Experimental and simulated locomotor phase duration characteristic. *Top*: The relationship between the locomotor phase and step cycle duration is shown with points representing the superimposed numerical and analytical solutions that are highly correlated with the experimental data lines; flexor (blue) and extensor (red) phases (Halbertsma, 1983). *Bottom*: The corresponding simulated speed (black points) is plotted as a function of step duration and compared to the experimental solution (black line) (see Fig. 3, in (Goslow et al., 1973)).

where *H* is the difference of simulated and experimental stance and swing periods. The experimental periods were calculated using a best-fit formula obtained empirically (Halbertsma, 1983). *M* is the difference of simulated and desired speed ranges that promotes the converging on nontrivial solutions. *O* is the cost associated with the erroneous co-activation of contralateral flexors. *C* is the degree of asymmetricity between the simulated speeds of the left and right limbs. All function components were normalized to the domain between 0 and 1 and relative weights (*k_1_, k_2_, k_3_, k_4_*)=(1, 0.7, 2, 0.4). *C* and *M* components were removed in simulations intended to produce asymmetrical gait (see Fig. 6 & 7 in Results).

### Optimization and parameter perturbation

Globally optimal sets of parameters were found numerically using a combination of the basin-hopping algorithm (Wales and Doye, 1997) in SciPy (Oliphant, 2007) and several constrained local minimizers: the non-linear optimization algorithm COBYLA (Powell, 1964), the truncated Newton algorithm (Nocedal and Wright, 2006), the L-BFGS-B algorithm (Byrd et al., 1995), and Powell’s method (Powell, 1964). *First*, the global optimal parameter set (*z*\*) was found. During optimization, the starting value for the basin-hopping algorithm was obtained from a brute force search over the complete parameter space. Other algorithms were then optimized sequentially to arrive at the optimal solution (*z*\* = *argmin(Jc)*). *Second*, we created a normal multivariate distribution to evaluate the nature of close-to-optimal solutions. For this, the distribution was defined by the mean at *z*\* and the covariance matrix with the diagonal elements set to 0.01*z*\* or the equivalent of the standard deviation set at 1% of the value of the optimal solution. The dataset of 10^5^ points was then drawn from this distribution and used in the comparison between the analytical and numerical solutions in Fig. 3A. *Third*, the intermediate solutions of the first step corresponding to local minima were selected to determine the full functional range of parameters in the model, excluding sets with large cost values (*Jc*>10). The adjusted for symmetricity range for each parameter is shown as the span of the y-axes in Fig. 4. *Fourth*, we used a uniform distribution across the symmetrical full range of parameters to create another dataset of 10^5^ values for the analysis of the expanded range comparison shown in Fig. 3B and 3C. *Fifth*, we created the parameter dataset perturbed by 10% from *z*\*. Similar to step 2 above, we created the normal multivariate distribution with the mean at *z*\* and the covariance diagonal elements set to 0.1*z*\*. *Sixth*, we randomly drew 40 starting seeds and tasked the basin-hopping algorithm (set to 200-iterations for each seed) to repeat the optimization using one of the four local optimization algorithms. This final step in the analysis generated 160 optimal sets for all local algorithms in our analysis. The comparison of parametric distributions is shown for a third of the best solutions in Fig. 4. The cut of solutions was necessary to reject expected minimization failures with non-converging searches or those terminating with large cost function values.

**Fig. 3.**
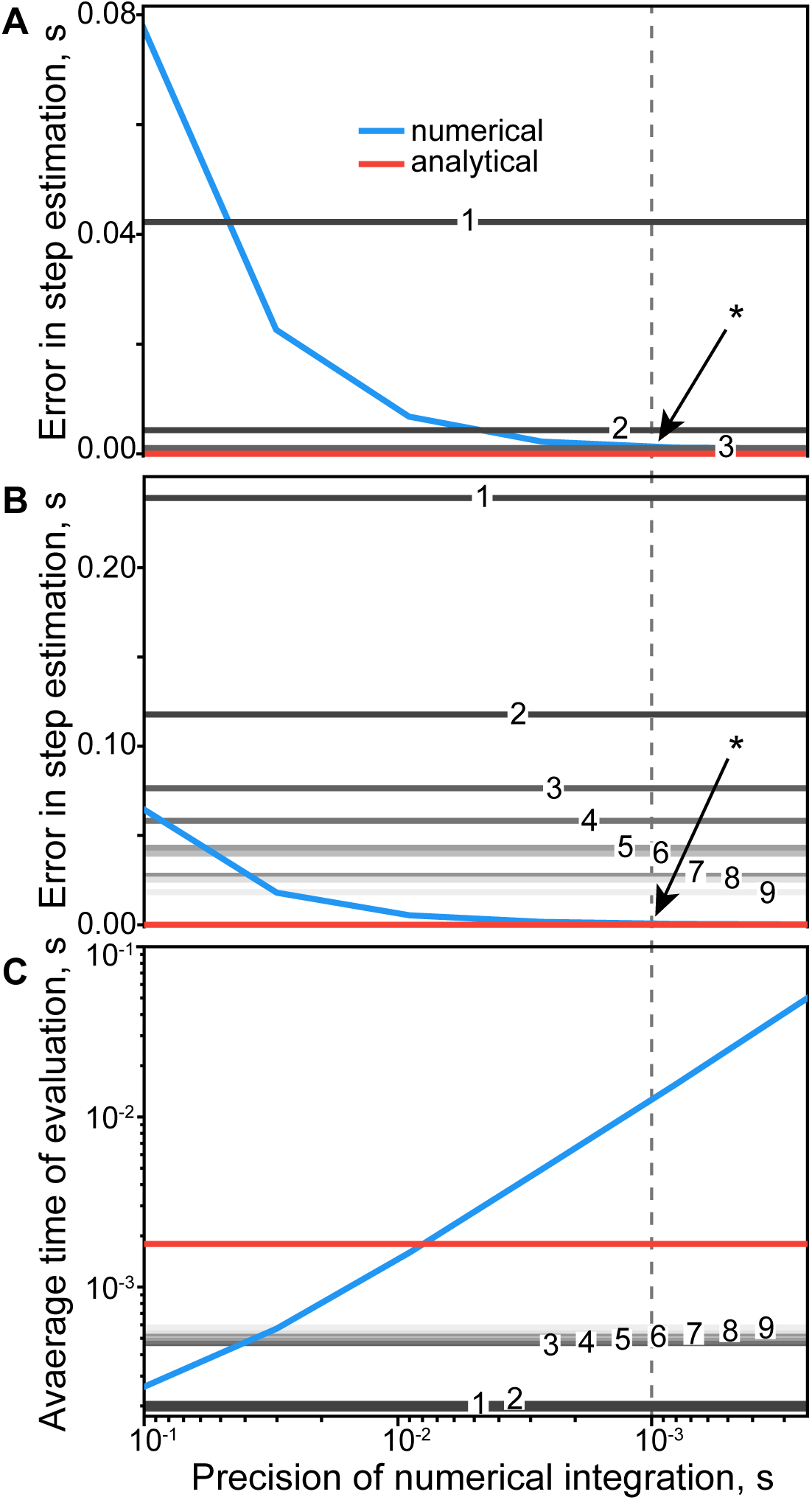
The comparison of analytical and numerical solutions. The measures of numerical (blue), analytical (red), and analytical approximations of different orders (shades of gray with order numbers) are plotted as functions of numerical precision, where the dashed line indicates the most relevant for real-time simulation precision of 1 ms. **A**. Full cycle error in the estimation of phase transition times using the 1% neighborhood of the optimal solution. Because the higher orders of approximations provide the same high precision as the cubic approximations, powers *τ*^4^-*τ*^9^ are not displayed. **B**. Similar to **A**, the errors are shown for the random distribution of parameters. **C**. Average CPU time needed to calculate a full step period of 1.25 s (average from Halbertsma’s equations) in Python/NumPy implementation. The data presented in all subplots was averaged over 10^5^ trials.

**Fig. 4.**
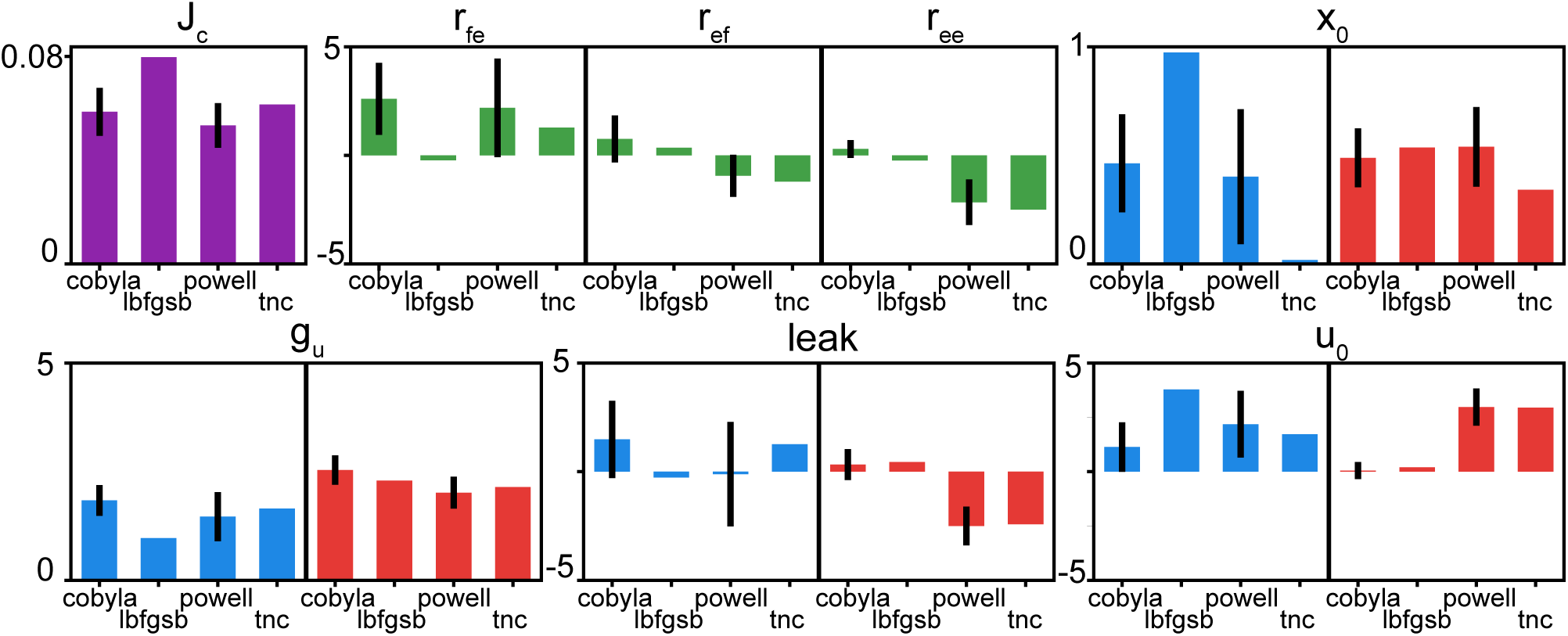
Analysis of parameter sensitivity. The distributions of model parameters and cost function (*Jc*) are shown for the selection of best optimization sets. Each subplot shows a mean with standard deviation of the parameter values in blue (flexor), red (extensor), and green (mixed) for 4 types of minimization algorithms. The vertical axis range reflects the full feasible range of parameters as determined by the examination of intermediate solutions (see step six in section “Optimization and parameter perturbation” of Methods), with the exception for the *Jc* values.

### Phenomenological models of locomotion

We used several phenomenological models created to describe the relationships between different parameters of stepping during locomotion in our analysis. The relationships between stance and swing phases relative to cycle duration were taken from the study by Halbertsma (1983)(Halbertsma, 1983). The relationship between step cycle duration (*T_c_*) and velocity (*V*) was taken from the study by Goslow et al. (1973), where *V*=*(1.84 · T_c_) ^1.68^* (see Fig. 2, bottom) (Goslow et al., 1973). Both studies used best-fit functions to describe data from a small sample of cats; yet, these relationships have been recently confirmed with a large subject pool (Frigon et al., 2015).

In the analysis of asymmetrical locomotion, we introduced a simple geometrical relationship for walking on a curve. The turn radius of an asymmetric bipedal walk (Eq. 8) was expressed as a function of hip width (*L*) and an asymmetry parameter *α*=*V_left_/V_right_*:

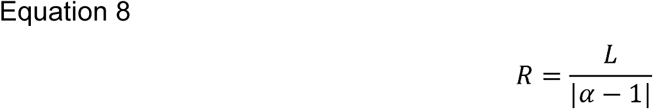

The corresponding heading direction change during a single step can then be stated as:

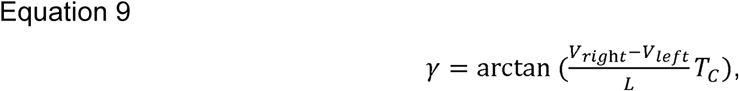

where denotes the heading direction angle from forward direction; *T_c_* - full step cycle period.

## Results

### Comparison of analytical and numerical solutions

In this study, the continuous dynamics between phase transitions was demonstrated with a simple CPG model expressed as a system of interacting oscillators and solved either numerically or analytically using an iterative algorithm (Eq. 4). Analytical solutions were validated in simulations producing experimentally observed periods of flexor and extensor activations in overground locomotion (for example, see Fig. 2). This model was further extended to analyze asymmetric gait and test the ability of this circuit to embed asymmetric gait control.

A high-precision numerical approach carries a processing cost that usually exceeds that of analytical methods. Fig. 3 shows the comparison of the processing cost between the numerical and analytical solutions for this model (Eq. 1). The error of evaluating phase transitions with the numerical method (blue line) and the analytical solutions using the root-finding algorithm (red line) was the same at the precision for numerical integration set to 10^-3^ s (intersection marked with *, Fig. 3A and 3B). The analytical solutions to Eq. 4, found by expanding the hyperbolic terms, linear to the 9th power, are shown with shades of gray in Fig. 3. Here, the difference between the analytical and numerical estimations of the time of phase transitions was evaluated with the root mean square metric of simulation quality. Shown in Fig. 3A, the quadratic approximation (gray line marked with a 2) provided similar quality to the analytical solutions (red line), with sets of close-to-optimal parameters (in 1% vicinity of the optimal set; see step two in section “Optimization and parameter perturbation” in Methods). When the model parameters were chosen randomly from the full range of feasible parameters (steps three and four in Methods), quadratic solutions did not provide desirable precision and performed worse than the numerical method, with other powers only approaching a reasonable threshold of over 10 ms error (Fig. 3B), which is the order of a motor unit action potential.

Fig. 3C shows that the analytical solution was the best choice for precise real-time applications of this model, outperforming the numerical method by close to an order magnitude. However, if estimation errors of over 10 ms are insignificant in a specific application, e.g. using EMG-driven simulations with aggressive low-pass filtering, then high orders of analytical approximations could provide appropriate solutions with even lower computational load than the full analytical solution. The approximations of powers 3-9 use the eigenvalue approach to find roots of polynomials, which is relatively costly but still more precise than some of the comparable numerical integrators.

### Parametric sensitivity

A perturbation analysis was used to investigate the parametric sensitivity of suboptimal solutions that satisfy Eq. 7. This analysis compared optimal values found by several different local minimization methods after a 10% normal parametric perturbation (for details, see steps five and six in section “Optimization and perturbation” of Methods). From 160 solutions, the 33% with the lowest *Jc* were: 30 by COBYLA, 1 by L-BFGS-B, 22 by Powell’s algorithm, and 2 by Truncated Newton’s. COBYLA and Powell’s algorithms provided 95% of the best solutions in this problem. The distribution of parameters in Fig. 4 with similar cost (*Jc*) across all methods indicates that similar outputs could be produced with disparate circuit parameters. The parameters in the model were differently conserved across similar solutions: the input weights (*G_u_*) had lower variability relative to other parameters, i.e. the static leak (*x_0_*), static input (*u_0_*), and interlimb connection weights (green, *r_ij_*).

### Behavioral implications of CPG morphology

The velocity hypothesis states that descending signals to a CPG are the desired speeds of each leg. We wanted to test further if the analytical solution to the ODEs would produce the same or a different velocity prediction for the modality of inputs. The direct relationship between the descending input and the temporal characteristics of stepping (step cycle, swing, and stance durations) was extracted from the second-order solution to Eq. 6. Although it has a complex non-linear form (Eq. 10), its combination with the solution from Goslow et al. (1973) for the relationship between step cycle period and forward velocity produced a linear result shown in Fig. 5 (*r*^2^=0.999, *p*<0.001 for left and right limbs).

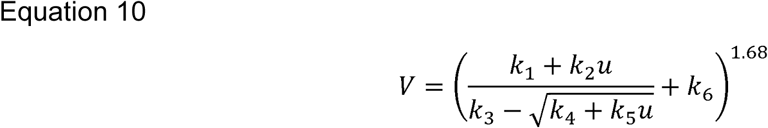

**Fig. 5.**
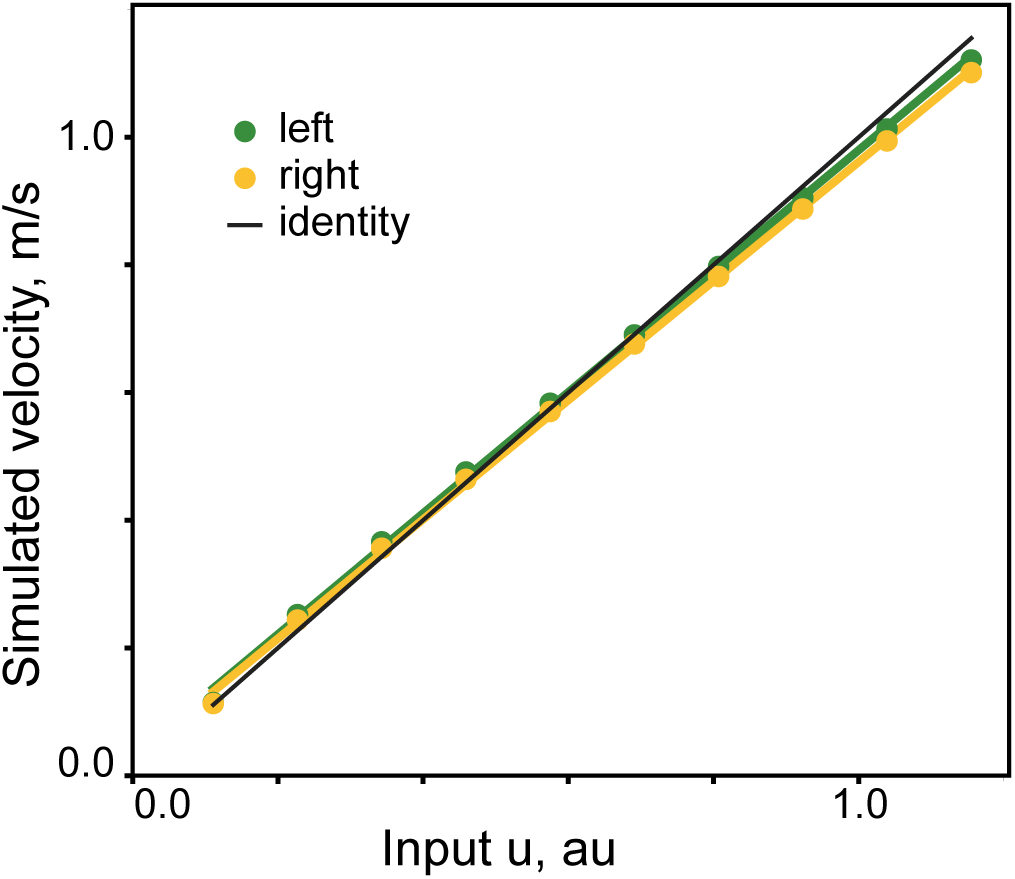
The relationship between the simulated CPG command signal to each limb and forward velocity. The analytical solution for the full step cycle was calculated over the set of 10 input values for each limb (*u*). Each value produced simulated step cycle duration values, which was then plotted as forward velocity calculated with the experimental relationship from Goslow et al. (1972) for each limb. The identity (*y*=*x*) is plotted in black.

where *k_i_* are configuration-dependent constants, *u* is descending input, and *V* is the forward velocity of locomotion.

We further explored the role of this descending command for velocity regulation in the generation of asymmetric gait. Asymmetric patterns were simulated by uncoupling the gains for the left and right inputs of both flexors and extensors _(_*g_uf1,_g_ue1,_g_uf2,_g_ue2_*_)_ in Eq. 2 and varying them independently by 33% of the optimal parameter set (Table 1). The C and M components responsible for pattern symmetricity and simulated velocity related errors were removed from the cost function (Eq.7) in this analysis. The simulated speed of walking for the left and right limbs was then calculated from the generated bilateral phases (Fig. 6). The parameter asymmetricity led to a steady gradient of the speed differences ( *α=V_left_/V_right_*, see Methods).

**Fig. 6.**
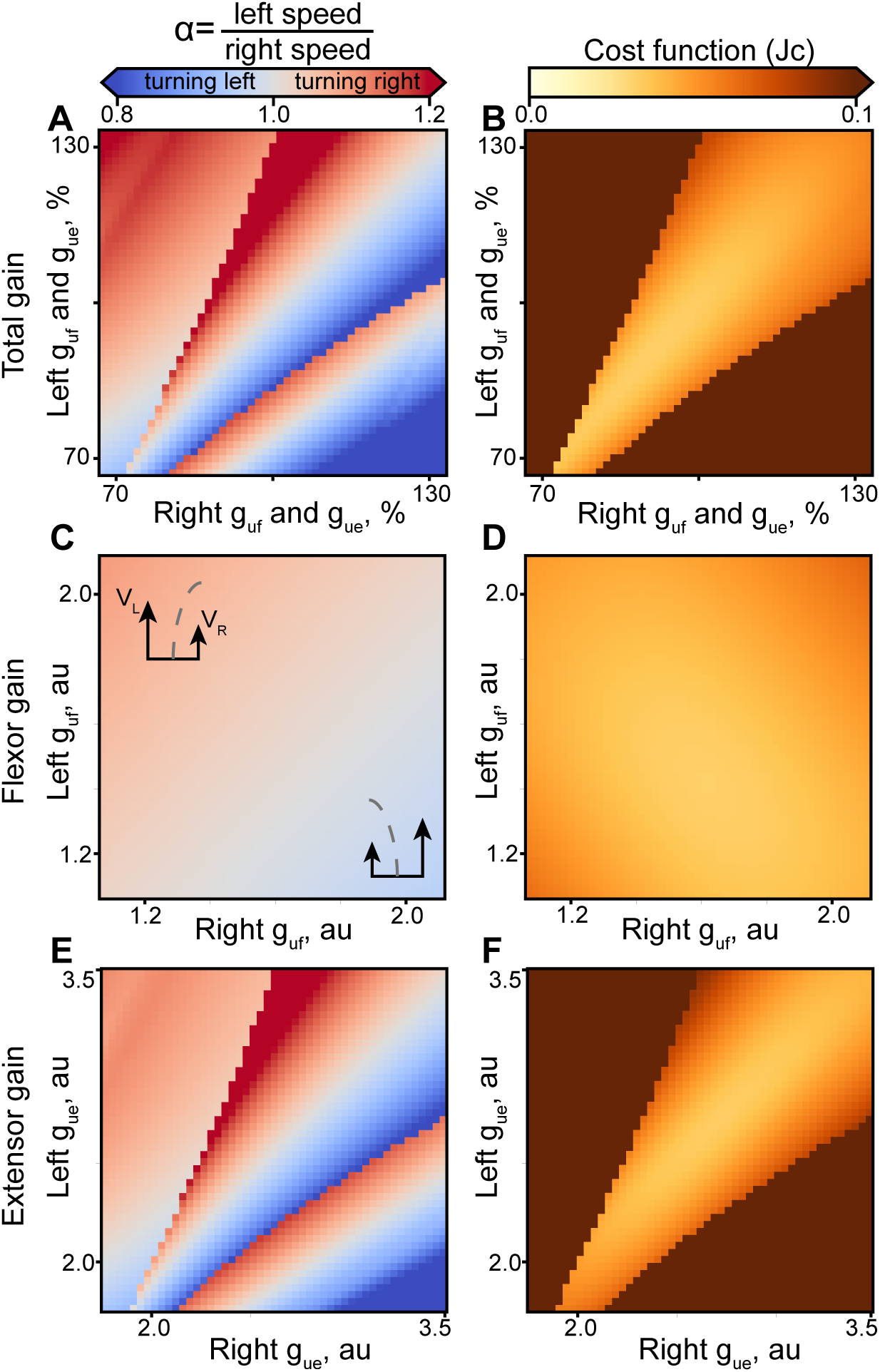
External inputs generate asymmetric gait in the model. The coupled and uncoupled input gain parameters (*g_ue_, g_uf_*) were related to the velocity asymmetry (*Left* panels) with the corresponding cost function outputs (*Right* panels). **A and B**. The input gains of flexors and extensors were varied together for each limb. **C and D**. Only flexor input gains (left and right *g_uf_*) were manipulated for each limb. **E and F**. Only extensor input gains (left and right *g_ue_*_)_ were manipulated for each limb. Inserts in **B** indicate the steering direction for two selected parameter sets.

**Table 1.** Optimal model parameters. The parameter set (*z*\*) for Eq. 2 that satisfies Eq. 7.

| $x_{0f}$ | $x_{0e}$ | $g_{uf}$ | $g_{ue}$ | $g_{lf}$ | $g_{le}$ | $u_{0f}$ | $u_{0e}$ | $r_{fe}$ | $r_{ef}$ | $r_{ee}$ |
| --- | --- | --- | --- | --- | --- | --- | --- | --- | --- | --- |
| 0.244 | 0.376 | 1.59 | 2.62 | -0.689 | 0.828 | 2.26 | -0.174 | -0.025 | 2.38 | 0.418 |

Figs 6A and 6B show that variation of both inputs (*g_uf_, g_ue_*) together can produce asymmetric walking, *α*=1.1, with the turn diameter as low as 10 m (calculated from Eq. 8, or heading direction *γ*=10° change per step, see Eq. 9). Only the parameter combinations corresponding to the continuous gradient around the midline produced appropriately accurate simulations with low *Jc* (Fig. 6B). Uncoupled inputs to flexors and extensors can similarly generate asymmetric gaits, with α up to 1.2 ( *γ*=20°). The gradient of cost for extensors was orthogonal to that for flexors in Figs 6D and 6F; the increased possible range of asymmetric speeds was associated with increased cost, as indicated in Fig. 6B, with the cost trough extending along the diagonal unity.

Fig. 7 shows that the intrinsic parameters in the model can also produce asymmetric gaits. Symmetric connections (e.g. in Eq. 2, *r_fe_*=*r_14_*=*r_41_*) were uncoupled (*r_14_*≠ *r_41_*) and varied independently. As in the analysis above, and *Jc* were calculated for parameter variations of up to ±33% of the optimal value. The connections from flexor to contralateral extensor did not provide a suitable gradient of asymmetric walking speeds in the explored range of parameters (Fig. 7A). Possible reasons are a low magnitude of the optimal value for this parameter (*r_ef_*, in Table 1) and the near constant relationship between swing duration and locomotor velocity (Fig. 2). The variation of extensor-to-flexor and extensor-to-extensor parameters (*r_ef_, r_ee_*_)_ produced asymmetric gaits (Fig. 7C and 7E) with a turn diameter of 10 m (heading direction *γ*=10° per step). These were comparable to the above result obtained from the analysis of external inputs. The profile of *Jc* was different for the gaits generated by variation of *r_ee_* and *r_ef_* parameters (Figs 7D and 7F). The extensor-to-flexor parameter *r_ef_* increased steering angle with a smaller increase in cost (Fig. 7F) than that of the extensor-to-extensor parameter, *r_ee_* (Fig. 7D). However, *r_ee_* could regulate asymmetric gaits over a larger range of velocities than *r_ef_*, as indicated by the diagonally extending trough in the cost function in Fig. 7F.

**Fig. 7.**
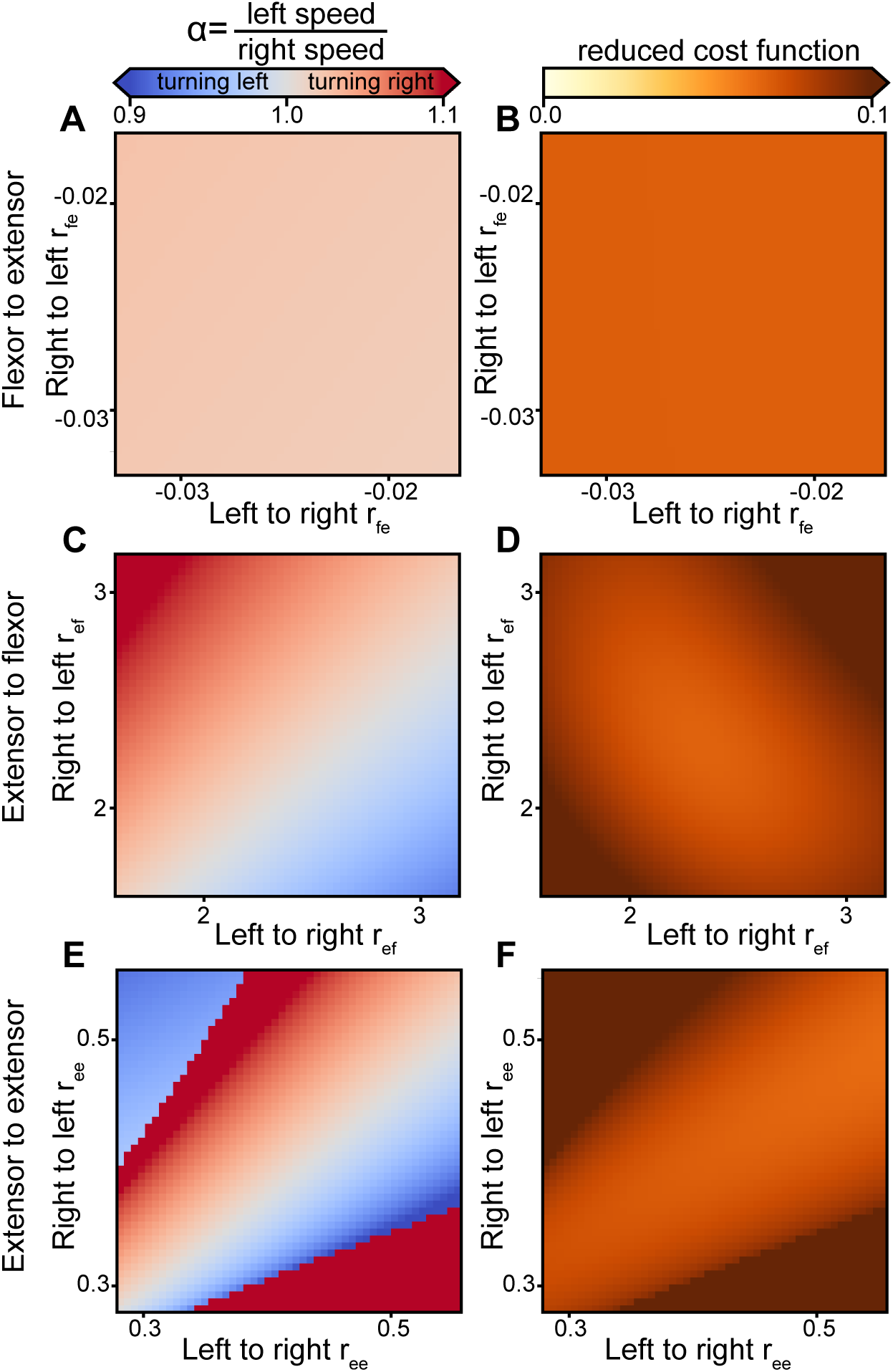
Intrinsic parameters generate asymmetric gait in the model. The uncoupled intrinsic parameters (*r_fe_, r_ef_, r_ee_*_)_ were related to the velocity asymmetry (*Left* panels) with the corresponding cost function outputs (*Right* panels). **A and B**. The flexor-to-extensor weights (*r_fe_*). **C and D**. The extensor-to-flexor weights (*r_ef_*). **E and F**. The extensor-to-extensor weights (*r_ee_*).

## Discussion

In this study, we developed a novel analytical description of a simple CPG model for locomotor phase timing and further expanded our previous model (Yakovenko et al., 2005) to include not only externally-driven asymmetric rhythmogenesis but also the opportunity to internalize this asymmetric transformation within the structure of a CPG. Our three central results are: i) the model can be solved analytically; ii) the analytical solution converges on the same conclusion that the input to the CPG is in the modality of limb forward velocity; and iii) the minimalistic model of a CPG built with coupled oscillating leaky integrators offers multiple opportunities for embedding asymmetric control.

### What is the goal of using analytical solutions of neurophysiological models?

Numerical solutions are usually the preferred option of solving complex models. For example, a biophysical CPG model proposed by Rybak et al. (2006) captures the neurological basis of activity in detail, often using hundreds of approximated parameters and their reconfiguration during failures in the motor execution (Rybak et al., 2006). Complex models with multiple estimated transformations may produce ensemble behavior that reproduces the expected outcome; however, the role of elements and their network properties are hard to predict and analyze. Unlike models that are not analytically solvable, simple models are often insightful and capable of identifying specific targets that modify circuit behavior (Tabak et al., 2000; Izhikevich, 2004; Barnett and Cymbalyuk, 2014b). For example, in the study of Barnett and Cymbalyuk (2014) the saddle-note bifurcation of equilibria was manipulated to design rhythmogenic regimes with appropriate timing and phase duration characteristics (Barnett and Cymbalyuk, 2014b). The employed bifurcation control method relies on the manipulation of a controlling parameter near a transition between different regimes responsible for spiking and bursting properties. Spardy et al. (2011) showed how the dynamical system analysis could identify the silent and bursting periods of system’s oscillation, the effect of sensory inputs on the range of behavior, and the operation of the CPG model in response to simulated spinal cord injury (Spardy et al., 2011a). This description was based on the simplified model (Markin et al., 2010; Spardy et al., 2011b) that uses two types of neuron implementations consisting of one- or two-dimensional differential equations for a single limb flexor-extensor CPG. Similar to other much more complex implementations, e.g. studies by Rybak et al. (2006), Morris and Lecar (1981), and Caplan et al. (2014) (Morris and Lecar, 1981; Rybak et al., 2006; Caplan et al., 2014), even this simplified formulation produces a challenging system of equations for 10 neurons with 33 connections between them. The model did noticeably have problems resolving locomotor phases for fast cycle durations (less than 800 ms, see Fig. 3 in (Spardy et al., 2011b)). In contrast, our simple model had only 4 parameters within a reciprocally connected system of 2 leaky integrators and simulated the same behavior without the aberrations at the extremes of experimental data (Yakovenko et al., 2005). This basic that we extended in this study was used to describe, for the first time, the novel flexibility of extensor- and flexor-dominant phase regulation.

As in other models, we were concerned that expanding the model’s parametric space to describe two limbs could introduce an uncontrollable increase in errors associated with the corresponding parametric expansion. The bilateral half-centers for two limbs required a system of 4 differential equations and the set of either 7 coupled (see Eq.2) or 16 uncoupled intrinsic and 4 extrinsic (input) parameters. The results for the expanded model in Fig. 2 showing phase modulation over the full range of walking velocities without limitations at the extremes was not a forgone conclusion. Overall, the increased parametric complexity in the model did not lead to an overfitting problem that could have appeared from estimating too many parameters from a low-dimensional set of behavioral data. Instead, the model consistently converged on similar solutions without the loss of validity indicated by the cost function.

Overfitting and underfitting are two major concerns in the selection of appropriate levels of abstraction for models (Lever et al., 2016). In the words of John von Neumann, “With four parameters I can fit an elephant and with five I can make him wiggle his trunk.” Here, our simple model relies on 20 parameters to generate low-dimensional output in the form of the phase characteristic in normal and asymmetric locomotion. Models based on Hodgkin-Huxley formalism could generate the same phase duration characteristic, albeit with the use of large model parameter sets that extend into hundreds and thousands. Remarkably, the solutions from these two different representations are similar, supporting the experimental and computational observations that the same network activity could be generated by the underlying disparate mechanisms (Prinz et al., 2004; Goaillard et al., 2009; Grashow et al., 2009; Caplan et al., 2014). Still, the convergence of our parameter search on the physiological network solution is validated only by the constraining behavioral data and extent of simulated validation using parameter sensitivity analysis. Even in this minimalistic model, the exploration of a 20-dimensional parameter space was challenging and led us to implement the analysis of a coupled symmetrical model first, where the parameters representing spinal neural elements mirrored across the midline were set to the same values. The perturbations in each parameter achieved with different minimization algorithms produced robust solutions, where small changes did not lead to large changes in outcome (Fig. 4). Thus, the model may not be overfitting for these particular phenomena under study.

### Embedding of asymmetric gait control in extrinsic and intrinsic parameters

Even in our relatively simple model, there is a complicated relationship between intrinsic connections and extrinsic inputs. An indication of this fact is the capacity for representing the same behavior within parameters corresponding to different anatomical structures. Thus, it was necessary to uncouple the parameters in Eq.2 to further extend the sensitivity analysis with the goal of exploring the functionality “hidden” in the complexity to generate falsifiable hypotheses or model predictions.

We chose asymmetric gait as the test task because it results from the normal control of steering or heading direction (Yakovenko, 2011; Galbreath et al., 2014), and it may contain indicators of long-term adaptations to injury. First, we “forced” the model to internalize the control of asymmetric stepping by changing only extrinsic parameters. The mechanism using only input gains of flexor half-centers, and less so extensor half-centers, was a robust method of changing the interlimb speed differential. This was also expressed as a change in the heading direction in this model. In Fig. 6, the tuning of input gains to flexor half-centers led to asymmetric speed ratios of 0.9 to 1.1, which corresponds to an estimated heading direction change of ±10° over one step cycle (about a 10 m turn diameter). This suggests that a single external input representing a heading direction could generate a realistic range of asymmetric gaits in this model. Second, we can similarly constrain the solution to the locus of intrinsic parameters responsible for the influences among four half-centers in the model. It was intriguing to see the capability of this model to embed the asymmetric processing within these pathways. Moreover, the simulations suggested that not all parameters are equal targets in that respect. The extensor-to-flexor and extensor-to-extensor (*r_ef_*, *r_ee_* in Fig. 7) parameters embedded the ability to generate asymmetric gaits with a reasonable turn diameter of 10 m, which is consistent with a “step turning” strategy, characterized by a wide base of support throughout the turn. It is likely that steeper turning would require the transition to a different “spin turning” strategy (Hase and Stein, 1999; Taylor et al., 2005). The alternative CPG configurations are illustrated in the schematic in Fig. 8. In studies of spinal segmental connectivity, these parameters would correspond to the ‘gains’ of propriospinal pathways connecting rhythmogenic networks within the spinal enlargement (Kiehn, 2011). Given the more rostral distribution of flexors than extensors within the lumbosacral enlargement (Yakovenko et al., 2002; Ivanenko et al., 2008) *r_ef_* and *r_ee_* pathways would have the network representations shown in Fig. 8B and C. Overall, relatively complex behavior, like steering, could be controlled with both extrinsic and intrinsic mechanisms available in this simple model.

**Fig. 8.**
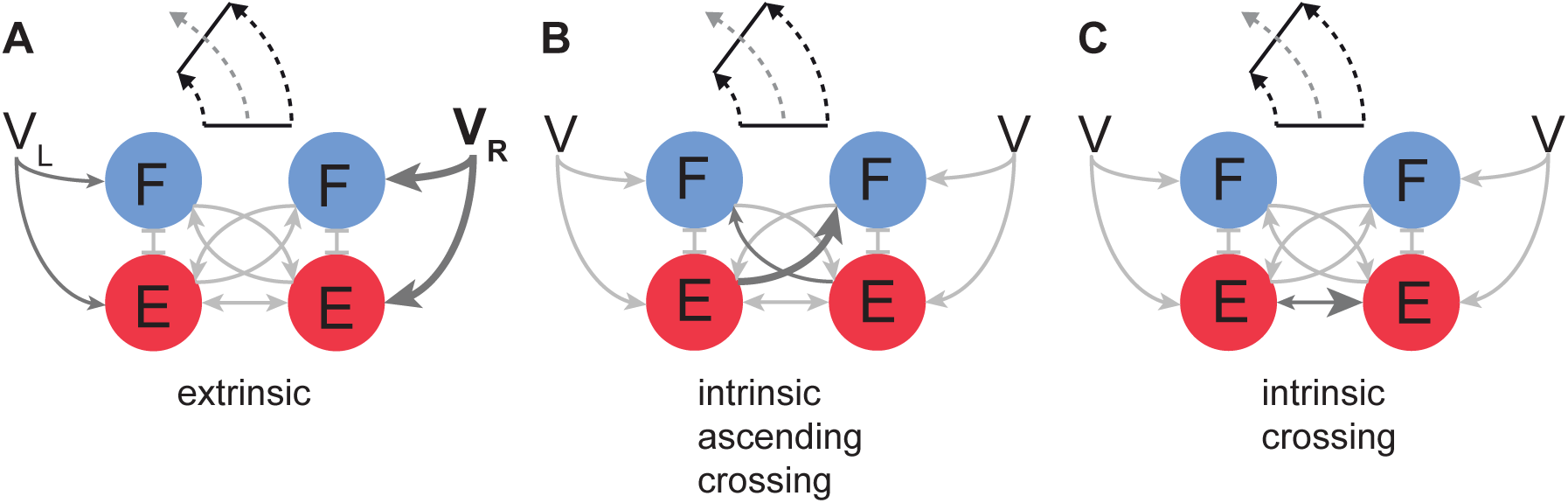
Schematic representation of multiple CPG configurations for steering. **A**. The configuration based on the external inputs to CPG. **B and C**. Two possible configurations of intrinsic connections producing the same asymmetric patterns as in **A**.

The analysis makes specific predictions about the propriospinal pathways that could be involved in long-term adaptations to asymmetricity. Human subjects could learn to compensate for the external perturbations applied to limbs while minimizing the overall limb impedance (Shadmehr and Mussa-Ivaldi, 1994; Dingwell et al., 2002). Even gross cortical inputs, like those generated by transcortical magnetic stimulation, can be compensated by the adaptation of transmission gains contributing to the regulation of locomotion (Schubert et al., 1999). Our results suggest that this adaptation can take place not only within pathways projecting to a CPG, but also within the limited locus of interactions between model’s half-centers. While this model has no realistic learning dynamics, the examination was limited to the naïve symmetrical and adapted asymmetrical states. This learning function could be implemented in future work with the use of simple learning mechanisms (Franklin et al., 2008; Wu et al., 2014) where intrinsic parameters could be updated under the reinforcement learning dynamics (Mahmoudi et al., 2013; Schultz, 2013). Overall, the model demonstrated that the general locomotor patterns for symmetric and asymmetric gaits may be achieved by the superposition of commands and intrinsic interactions within the minimalistic structure of a CPG. This novel flexibility of functional representation for asymmetric pattern generation has not been previously demonstrated in models, and it posits specific predictions for mal- or adaptations to asymmetry due to peripheral or central abnormalities.

### The simple model of locomotor rhythm generation

This model is not likely producing the overfitting of behavior as indicated by the sensitivity analysis. Still, there is the possibility that this model is instead underfitting the locomotor patterns associated with asymmetric gait. To discuss the appropriate level of abstraction that limits the possibility of underfitting for this task, we need to examine the concept of neuromechanical tuning (Prochazka and Yakovenko, 2007; Ting et al., 2015). Specifically, locomotor control is a phenomenon produced by multiple elements that combine predictive and reactive functions. In analogy with the Smith’s predictor (Smith, 1957), the specific role of the CPG is to predict the mechanical interactions between the limb and ground. To this extent, our model can reproduce the transformation from input speeds to appropriate inter- and intra-limb coordination of multiple muscle groups without the need for molecular level dynamics (Yakovenko, 2011). The CPG function could then be specified as a dynamical transformation of simple high-level signals into complex granular functional subdivisions of temporal activations appropriate for locomotion. Both analytical and numerical solutions of our minimalistic CPG model support the hypothesis that the main function of a CPG is the transformation of high-level locomotor signals associated with whole limb function, i.e. the speed of locomotion, into low-level phasic activity patterns of limb muscles. This computational inference agrees with previous studies demonstrating that the one-dimensional input to the MLR in the form of stimulation magnitude or frequency can be transformed by a CPG into specific velocity-dependent phasic activity in vertebrates (Shik et al., 1966; Smetana et al., 2010). Thus, the underfitting for CPG models describing the phase duration characteristic would be classified by the inability to use high-level signals related to the forward velocity as the control signal for asymmetric gait. We demonstrated that this model can readily transform limb velocity-related inputs into asymmetric phase characteristics. Moreover, the model can embed these high-level representations within its internal structure. As shown previously (Yakovenko et al., 2005), it can also generate both flexor-dominated and extensor-dominated phase regulation at different speeds.

To conclude, in this paper we report for the first time a model of bilateral CPG with analytical and numerical solutions capable of generating symmetrical and asymmetrical gaits appropriate for whole body steering. The steering behavior can be generated by either extrinsic limb velocity related inputs to left and right half-center oscillators or embedded asymmetry within intrinsic propriospinal gains from extensor half-centers to the contralateral flexor or extensor half-centers. Moreover, these asymmetric changes may correspond to either a natural control of limb velocity adjustments regulating the heading direction or pathological changes to the inputs or structure of the locomotor CPG. The existence of multiple network states capable of generating the same empirical observations is a novel identified challenge for CPG models.

## Acknowledgements

The authors thank Dr. Jonathan Rubin for the general discussion of analytical solutions in locomotor CPG modeling. We thank the West Virginia Clinical and Translational Science Institute for editorial support.

